# Extreme C-to-A Hypermutation at a Site of Cytosine-N4 Methylation

**DOI:** 10.1101/2021.01.19.427316

**Authors:** Joshua L. Cherry

## Abstract

Methylation of cytosine in DNA at position C5 increases the rate of C→T mutations in bacteria and eukaryotes. Methylation at the N4 position, employed by some restriction-modification systems, is not known to increase the mutation rate. Here I report that a *Salmonella enterica* Type III restriction-modification system that includes a cytosine-N4 methyltransferase causes an enormous increase in the rate of mutation of the methylated cytosines, which occur at the overlined C in the motif 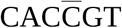. Mutations consist mainly of C→A transversions, the rate of which is increased ~500-fold by the restriction-modification system. The rate of C→T transitions is also increased, and somewhat exceeds that at C5-methylated cytosines in *Dcm* sites. Two other *Salmonella* N4 methyltransferases investigated do not have such dramatic effects, although in one case there is a modest increase in C→A mutations along with an increase in C→T mutations. Sensitivity of the C→A rate to orientation with respect to both DNA replication and transcription is higher at hypermutable sites than at other cytosines, suggesting a fundamental mechanistic difference between hypermutation and ordinary mutation.

**IMPORTANCE:** Mutation produces the raw material for adaptive evolution, but also imposes a burden because most mutations are deleterious. The rate of mutation at a particular site is affected by a variety of factors. In both prokaryotes and eukaryotes, methylation of C at the C5 position, a naturally-occurring DNA modification, greatly increases the rate of C→T mutation. A distinct C modification that occurs in prokaryotes, methylation at N4, is not known to increase mutation rate. Here I report that a bacterial restriction-modification system, found in some *Salmonella*, increases the rate of C→A mutation by a factor of 500 at sites that it methylates at N4. This rate increase is much greater than that caused by C5 methylation. Although fewer than one in 1600 positions analyzed are methylation sites, over 10% of all mutations occur at these sites. Like other examples of extremely high mutation rate, whether naturally-occurring or the result of laboratory mutation, this phenomenon may shed light on the mechanism of mutation in general.

## OBSERVATION

Methylation of cytosine in DNA at the C5 position increases the rate of C->T transition in bacteria (1, 2) and eukaryotes (3, 4). Some restriction-modification (RM) systems employ a second type of cytosine modification, methylation of the exocyclic N4. This modification is chemically analogous to the more common methylation of adenine at the exocyclic N6. As far as is known, cytosine-N4 methylation does not increase the rate of mutation.

Comparison of genome sequences of closely-related bacteria allows estimation of mutation rates in different sequence contexts. The NCBI Pathogens database provides variation information and phylogenetic trees for clusters of closely-related bacteria. Hypermutation due to cytosine-C5 methylation was detectable from this data (2).

### Hypermutation at 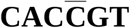 in a *Salmonella* cluster

Investigation of nearest-neighbor effects on mutation rate in *Salmonella enterica* revealed an atypically high rate of CCG→CAG mutation (including CGG→CTG mutation on the opposite strand) in a cluster of closely-related isolates of serovar Typhimurium (NCBI Pathogens cluster identifier PDS000026710.24) (Fig. S1). This cluster contains 1274 isolates.

Further investigation (Fig. S2) revealed that the high rate of C→A mutation is specific to the hexamer 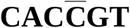, the hypermutation occurring at the third, overlined C. Of the 1228 CCG→CAG changes in the chromosomal coding sequences, 1104 occur at this motif. These account for the excess at CCG, and constitute 10.5% of all mutations. Several types of evidence indicate that this phenomenon is not an artifact of sequencing errors (Text S1).

The occurrence of such a large fraction of mutations in a hexamer motif, which occurs only 2582 times in the 4.27Mb of sequence analyzed, indicates an extraordinarily high mutation rate. The acceleration at these sites can be calculated by comparison with the other *Salmonella*, with the exclusion of other clusters exhibiting the phenomenon (see below). The rate of C→A mutations at this motif in this cluster is ~500 times that in others. By comparison, the rate of transitions at C5-methylated cytosines ranged from ~8 to ~50 times the rate at non-methylated cytosines (2).

The rate of C→T mutations is also elevated at 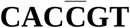. This effect is more modest than the increase in C→A mutation, but nonetheless fairly strong: the rate of C→T changes at these sites is 10.6 times higher at in the cluster than in those unaffected by hypermutation. The transition rate at these sites is somewhat higher than that at *Dcm* hotspots.

### Hypermutable cytosines are positions of N 4 methylation by a restriction/modification system

Among the DNA methyltransferases encoded by the genome of the reference isolate for this cluster, CFSAN001921 (RefSeq accession NC_021814.1), is the modification subunit of a Type III restriction/modification (RM) system (RefSeq accession WP_020837120), which is present almost universally in the cluster. A blasp search of the REBASE database (5) found several exact matches to the methyltransferase in *Salmonella*. PacBio sequence data, which allows determination of methylation motifs (6, 7), i s available for three of the genomes. Results for strain FDAARGOS_312 report cytosine N4 methylation at the overlined residue in 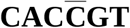, precisely the motif at which hypermutation occurs. REBASE remarks that methylation is likely due to this methyltransferase because there are no other plausible candidates. Thus, this Type III restriction/modification system appears to be responsible for the hypermutation.

This system is encoded by an element integrated at the *ssrA* transfer-messenger RNA locus. The element encodes an integrase and carries other genes typically associated with mobile elements. The RM system may have entered *Salmonella* recently, perhaps from a host in which it does not cause extreme hypermutation.

Several smaller clusters also contain the methyltransferase in most or all isolates. As shown in Fig. 1, these also display an increased rate of C→A mutation at 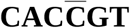 sites. These clusters are, according to the tree provided in the NCBI Pathogens data, all closely related to PDS000026710.24. Although they do not form a clade in that tree, they might do so in reality.

**FIG 1.**
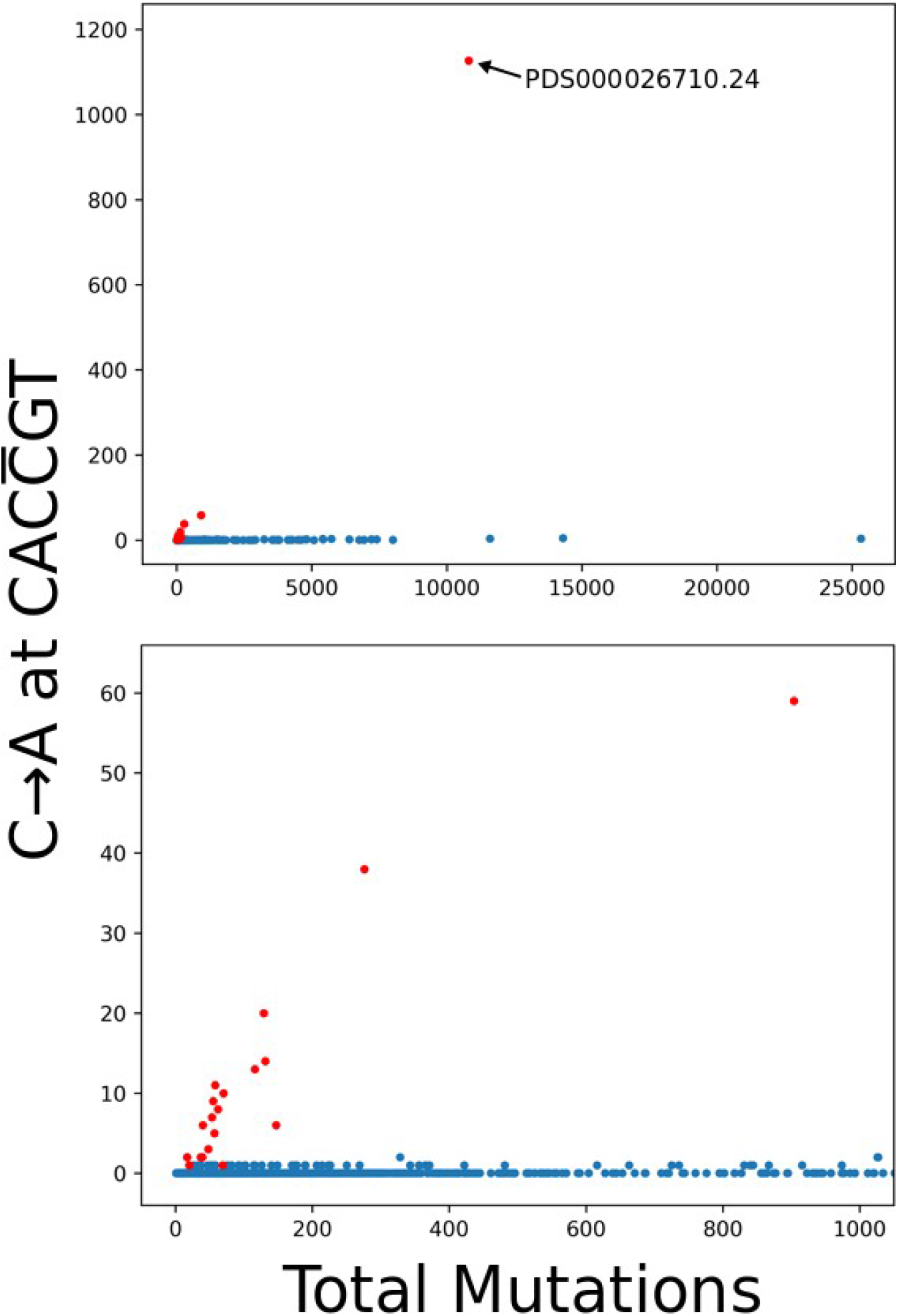
Number of C→A mutations at hypermutation sites *vs*. total mutations for each *Salmonella* cluster. Red points represent clusters in which most or all isolates carry the Type III RM system that is apparently responsible for hypermutation. The lower panel is a zoom of the upper panel.

Independent confirmation that the RM system causes hypermutation comes from a distantly-related cluster in which a single isolate contains it. There are two clusters in which a minority of isolates carry the system. In one, two isolates that descend directly from the same multifurcating node (PDT000310850.2 and PDT000315698.1 in cluster PDS000026701.8) carry it, but none of the ten mutations in the relevant branches are at 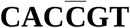. This result is not inconsistent with hypermutation, since the expected number among ten mutations is only slightly more than one. In the more informative case, a single isolate in the cluster carries the RM system (PDT000338580.1 in PDS000028569.6). Two of the eleven mutations on the terminal branch leading to this isolate occur at 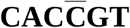, one a change to A and the other a change to T. This would be extremely unlikely without hypermutation. No other changes at this motif occur in the cluster.

### Effects of other cytosine-N4 methyltransferases

REBASE lists two other groups of cytosine-N4 methyltransferases. *Salmonella* isolates carrying the M.StyI or close relatives, which methylate at CCWWGG, show no sign of hypermutation to A or T. This may be due to lack of power: the 95% confidence interval for the relative rate to A is 0.056-13.2 (Fisher’s exact test), though for T it is 0-2.3. In isolates carrying M.SptAI or close relatives, which methylate at CAGCTG, the rate of mutation to A at these sites is increased by a factor of 7.2 (95% confidence interval 3.2-14.7), and to T by a factor of 8.6 (6.0-12.3). These increases are substantial and noteworthy, but do not approach the ~500-fold increase caused by the Type III system.

### Clues to mechanism

Extreme hypermutation does not appear to be a general consequence of cytosine-N4 methylation. Thus, some other aspect of the Type III system likely plays a role in hypermutation. Some candidates are the sequence context of the modified C, an activity of the restriction endonuclease (e.g., nicking), and a mutagenic side-reaction of the methyltransferase.

A possibility in the last category is transfer of a methyl group to an atom other than N4, such as N3 of the target cytosine. Methylation of N3 has been observed as a side-reaction of a cytosine-C5 methyltransferase (8). This modification is mutagenic in *E. coli* when present at one position in a singled-stranded DNA (9). In a wild-type background it leads mainly to C→T mutations, but the outcome might be different in the chromosome because is double-stranded and possibly N3-methylated at many positions. The outcome of replication of N3-methylcytosine varies among human DNA polymerases, and is in some cases incorporation of a T (10), which potentially leads to C→A mutation.

The rate of C→A mutation at hypermutation sites is affected by orientation with respect to both the direction of chromosome replication and the direction of transcription. It is higher by a factor of 4.5 when the C, rather than the paired G, is on the leading strand. At other sites, the C→A rate is only 1.3-fold higher on the leading strand. The rate at hypermutation sites is higher by a factor of 1.5 when the C is on the non-template strand for transcription. For other sites this factor is 1.1 and is statistically indistinguishable from unity. These contrasts likely reflect mechanistic differences between mutation at hypermutation sites and ordinary mutation.

The most common mechanism of C:G→A:T transversions involves 8-oxoguanine, a product of oxidative DNA damage (11). The hypermutation reported here might involve increased formation or incorporation of 8-oxoguanine opposite the affected cytosine, or an increased probability that it leads to mutation. If so, mutational inactivation of protections against 8-oxoguanine (11) might have a synergistic effect on hypermutation.

## ACKNOWLEDGMENTS

This research was supported by the Intramural Research Program of the NIH, National Library of Medicine. The findings and conclusions in this report are those of the author and do not necessarily represent the official position of the U.S. National Institutes of Health or Department of Health and Human Services.

