## Supplementary Figs. S1 and S2 and Text S1 for "Extreme C-to-A Hypermutation at a Site of Cytosine-N4 Methylation"

for

Joshua L. Cherry

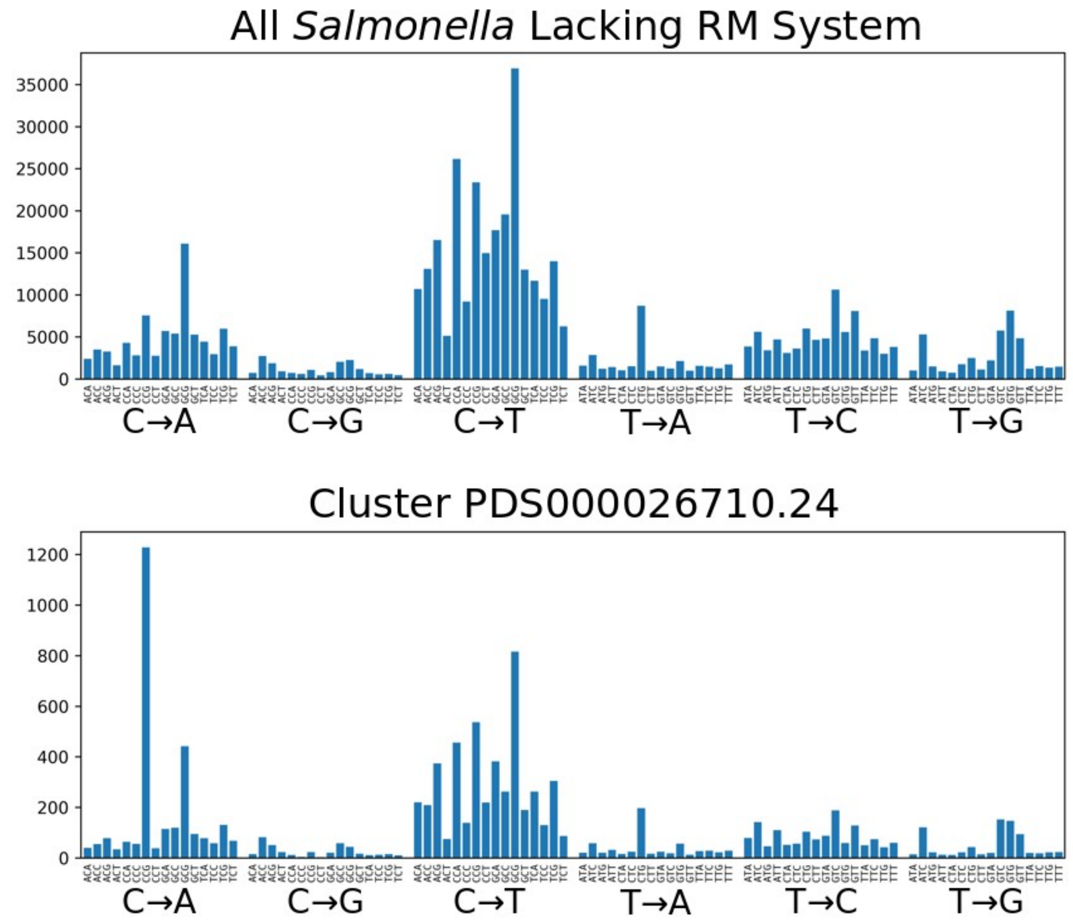


**FIG S1** Counts of mutations of all six strand-symmetrized types in all 16 possible nearest-neighbor contexts. A large comparative excess of C→A mutations in the context CCG for is obvious in the lower panel, which represents the cluster of isolates in which hypermutation was first detected.

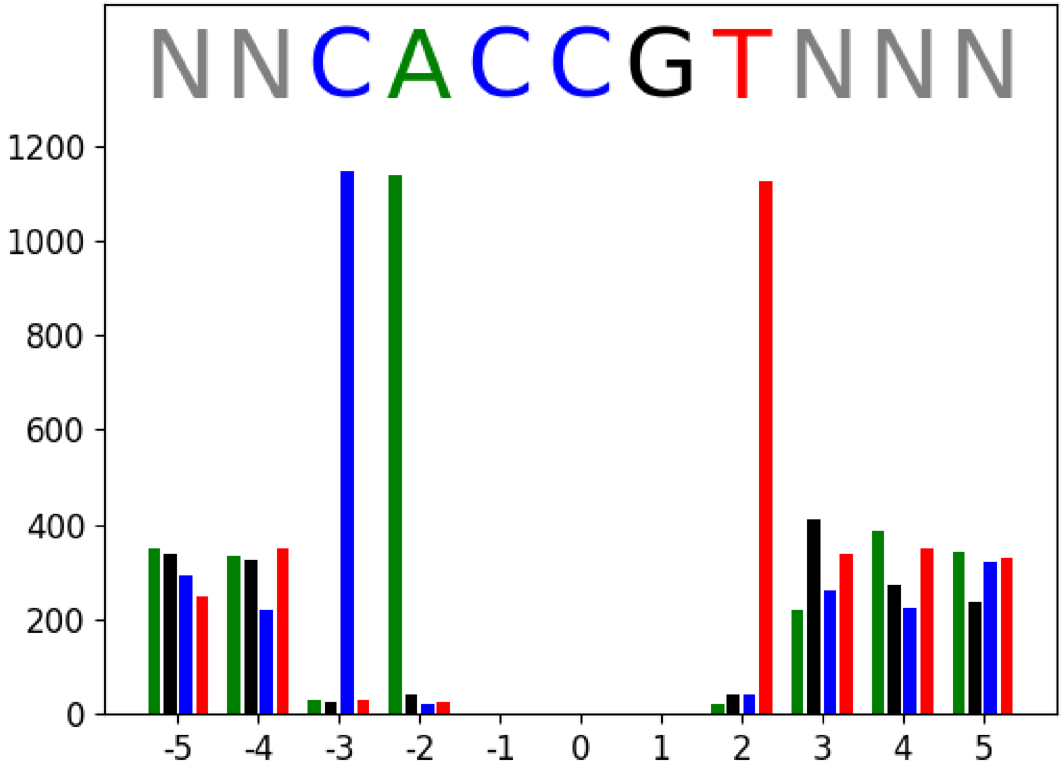


**FIG S2** Nucleotide frequencies at positions surrounding CCG in cases of mutation to CAG in cluster PDS000026710.24. Counts of A, G, C, and T are represented by green, black, blue, and red bars, respectively. At three surrounding positions, one nucleotide predominates, revealing that the excess mutations at CCG occur more specifically at CACCGT.

### Text S1

#### Ruling out Sequencing Error

The apparent hypermutation reported here cannot be the result of a general systematic sequencing error, since it only affects a few clusters and appears to depend on an RM system whose methyltransferase recognizes the hypermutation motif. However, the possibility of systematic error caused by the cytosine N4 methylation, or some other activity of the RM system, must be considered.

Errors of this type seem unlikely, as they would seemingly require an implausibly high error rate in the replication of the modified template in PCR (part of Illumina sequencing, which was used for most isolates). In addition to this consideration, several types of evidence rule out the possibility that the effect is due to sequencing errors:

1. Sequence changes that occur on internal branches must be natural mutations. Although the high polymorphism at the hexamer motif increases the chance of erroneous mapping to an internal branch, consideration of internal branches is enlightening. Of the inferred C→A changes in the cluster of interest, 30.9% are on internal branches (including implied resolution of multifurcations). This fraction is nearly the same as that for other mutations in the cluster, 30.1%. There is no obvious reason that these fractions would be so similar if sequencing errors caused the apparent hypermutation. The fraction on internal branches would likely be much lower in that case.

2. The presumptive mutations contain much phylogenetic signal that agrees with the tree of the isolates, even if the tree is inferred without the use of apparent hypermutation sites. There are 137 instances of the hexamer motif at which exactly two isolates are affected by a C→A change. In 59 of these, or 43%, the two isolates are sisters (descend from the same node) . The fraction expected for sequencing errors that occur uniformly and independently among isolates is 0.14%. If there are batch effects on sequencing errors, and batches tend to contain closely-related isolates, sister pairs will be represented more highly. However, 26 of the sister pairs come from different sequencing centers. These pairs alone, which cannot be due to batch effects, greatly exceed the expectated number for sequencing errors.

3. The terminal branch length index (TBLI) (S1) is based on the fact that naturally-occurring mutations occur disproportionately on long branches. Although originally applied to detection of laboratory mutation, it can also detect sequencing errors, which are similarly expected to occur without regard to terminal branch length. The TBLI for C→A mutations at the hexamer motif in cluster PDS000026710.24 is 0.984, very close to the value of one expected for natural mutations. A value of zero, expected for sequencing errors, can easily be rejected statistically ($p$=9.6e-74).

4. If the apparent C→A mutations at the hexamer motif were due to sequencing errors, the sequence reads for affected isolates would likely contain many reads bearing the original C at affected positions. Similarly, isolates for which the assembled sequence had a C at the position would likely

contain many reads with an A. According to the counts of mapped reads in the VCF file (NCBI Pathogens data), this is not the case. Few conflicting reads occur at such positions for either type of isolate.

5. The observed dependence of the mutation rate on orientation with respect to replication and transcription would not be expected for sequencing errors.

6. Most of the sequences in the cluster were determined by Illumina sequencing, but two were determined by PacBio SMRT sequencing. One of these has an A rather than the more frequent C at two sites with the hexamer context, in agreement with its neighbors in the phylogenetic tree. Detection of such differences by two different sequencing technologies adds to the evidence that they are genuine.
